# Emergence of multiple set-points of cellular homeostatic tension

**DOI:** 10.1101/2022.07.02.498579

**Authors:** Yuika Ueda, Shinji Deguchi

## Abstract

Stress fibers (SFs), a contractile actin bundle in nonmuscle mesenchymal cells, are known to intrinsically sustain a constant level of tension or tensional stress, a process called cellular tensional homeostasis. Malfunction in this homeostatic process has been implicated in many diseases such atherosclerosis, but its mechanisms remain incompletely understood. Interestingly, the homeostatic stress in individual SFs is altered upon recruitment of α-smooth muscle actin in particular cellular contexts to reinforce the preexisting SFs. While this transition of the set-point stress is somewhat a universal process observed across different cell types, no clear explanation has been provided as to why cells end up possessing different stable stresses. To address the underlying physics, here we describe that imposing a realistic assumption on the nature of SFs yields the presence of multiple set-points of the homeostatic stress, which transition among them depending on the magnitude of the cellular tension. We derive non-dimensional parameters that characterize the extent of the transition and predict that SFs tend to acquire secondary stable stresses if they are subject to as large a change in stiffness as possible or to as immediate a transition as possible upon increasing the tension. This is a minimal and simple explanation, but given the frequent emergence of force-dependent transformation of various subcellular structures in addition to that of SFs, the theoretical concept presented here would offer an essential guide to addressing potential common mechanisms governing complicated cellular mechanobiological responses.

## 1. Introduction

Stress fibers (SFs), a contractile actin bundle, are known to intrinsically sustain a constant level of tension in nonmuscle mesenchymal cells (Kaunas and Deguchi, 2011). Specifically, the force sustained in individual SFs ranges from approximately one to several tens of nano-Newtons, while their cross-sectional area is several square micrometers, giving rise to a tension or tensional “stress” (i.e., the force divided by the cross-sectional area) of several nN/μm^2^ (Balaban et al., 2001; Tan et al., 2003). Under this cellular regulation called tensional homeostasis, perturbations applied to SFs upon mechanical stresses such as cyclic stretch or fluid shear stress onto the cells are relaxed over time through structural remodeling, allowing the stress in SFs to remain at the intrinsic set-point value (Kaunas et al., 2006). Malfunction in this homeostatic control or adaptive response has been implicated in chronic pro-inflammatory states that potentially lead to diseases such as atherosclerosis and cancer (Chien, 2007; Hahn and Schwartz, 2009).

Interestingly, the set-point stress in SFs is altered upon recruitment of α-smooth muscle actin (α-SMA) (Hinz et al., 2001). Specifically, Hinz and colleagues found in myofibroblasts that a homeostatic stress of ∼3.8 nN/μm^2^ is sustained at individual focal adhesions (FAs), to which SFs are serially connected at their termini, with an area of ≦7.5 μm^2^; but, at other FAs with a larger area of >7.5 μm^2^ to which they refer as “supermature” FAs, a greater magnitude of stress of ∼12 nN/μm^2^ is instead sustained (Goffin et al., 2006; Hinz, 2010). This jump in the set-point stress value is induced by modulating mechanical cues to increase the magnitude of the force in the connected SFs, for example by plating cells on a stiff substrate, and is accompanied by new association of α-SMA to the preexisting SFs, with which originally β-cytoplasmic actin isoforms are associated. Supermature adhesions are induced in other cell types such as osteoblasts and hepatic stellate cells (Biggs et al., 2008; Liu et al., 2010). Similar force-dependent transformation of small adhesions into large ones called fibrillar adhesions have also been reported in endothelial cells and smooth muscle cells (Zaidel-Bar et al., 2003; Gardel et al., 2010; Saito et al., 2014). Collectively, the force-driven transition of the homeostatic cellular stress that individual SFs sustain is thus a somewhat universal process.

Despite these observations, no clear explanation has been provided as to why cells end up possessing different stable stresses. In the present theoretical work, we describe that a specific constitutive model depicting the nature of SFs yields the presence of multiple set-points of the homeostatic stress, which transition among them depending on the magnitude of the sustained force. This is a simple explanation, but given the frequent emergence of force-dependent transformation of various subcellular structures in addition to that of the SF– FA complex, the concept presented here may provide a fundamental insight into the underlying physics of complicated cellular mechanobiological responses.

## 2. Methods

### 2.1 Model description

α-SMA is incorporated to and reinforces preexisting SFs when they are allowed to exhibit a contractile force that is high in magnitude enough to trigger the incorporation (Hinz et al., 2001, 2010; Goffin et al. 2006) (Fig. 1a). It has also been reported that cellular elastic modulus is elevated upon increased localization of α-SMA along the length of SFs (Liu et al., 2013; Sakamoto et al., 2016; Liu et al., 2018; Stylianou et al., 2018; Wang et al., 2019). Given these observations, we assume that the elastic modulus of individual SFs *E* is altered as a function of the contractile force *F*, produced and sustained in the SFs, in a sigmoid manner described by

**Fig. 1.**
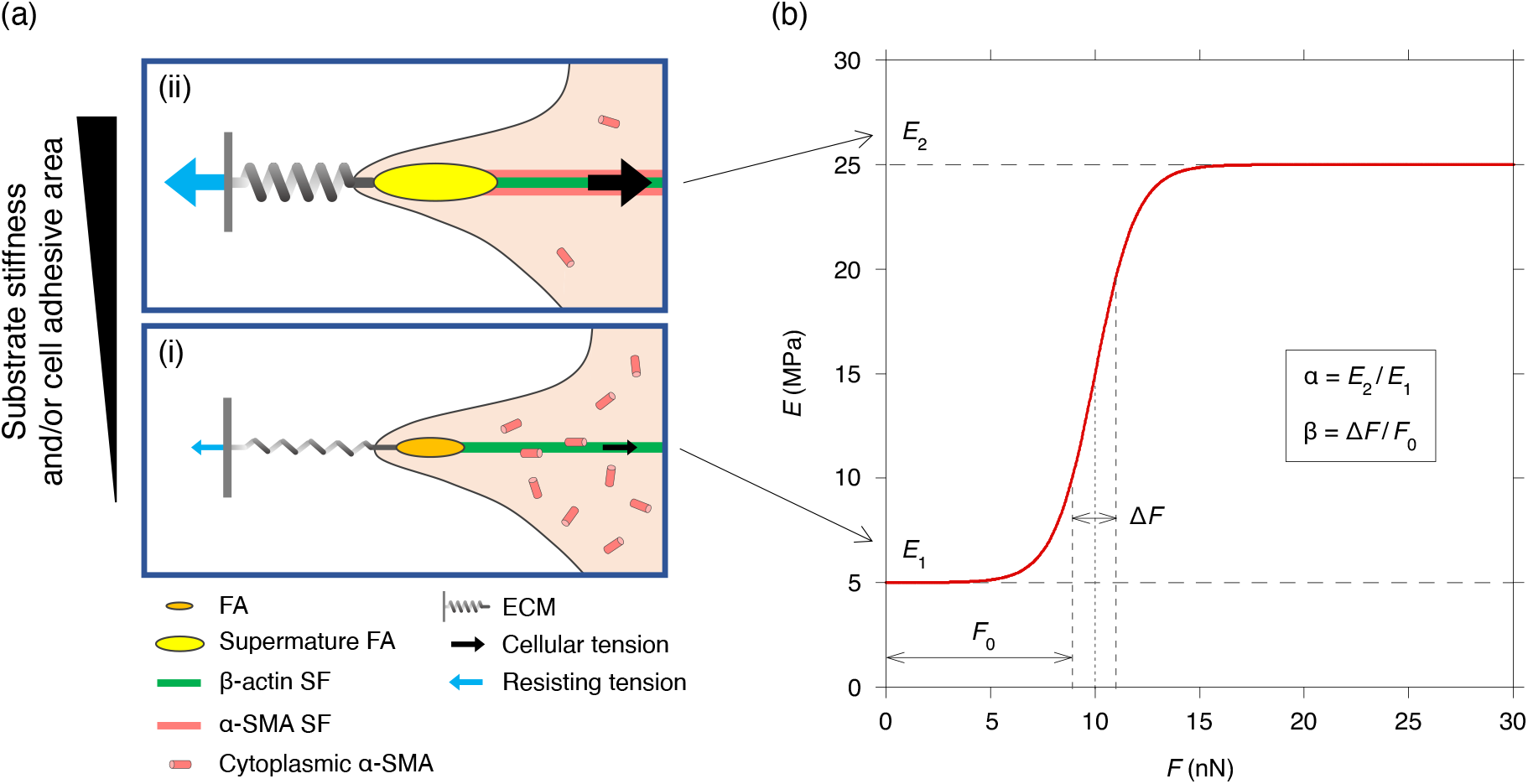
Force-dependent increase in elastic modulus of SFs. (a) α-SMA incorporation to a large tension-bearing SF depicted according to Goffin et al. (2006). Tension is relatively low (i) and high (ii) in magnitude at a soft and stiff extracellular milieu, respectively, as represented by the elongation of the extracellular matrix (ECM) spring. (b) The constitutive relationship for the elastic modulus of SFs *E* as a function of the tension *F* described by Eq. (1).

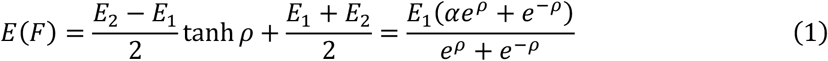

Where

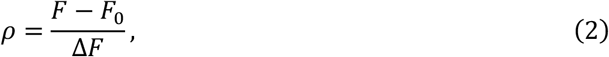

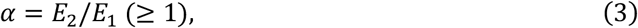

and *E*_1_ *E*_2_, *F*_0_, and Δ*F* denote the lower and upper limits of the elastic modulus, the force that provides the average elastic modulus, and the characteristic range of force where this transition takes place, respectively (Fig. 1b). More specifically, *E*_1_ and *E*_2_ represent the elastic modulus of SFs without (low-force) and with (high-force) α-SMA incorporation, respectively. The rationale for using the sigmoid function comes from the fact that the elastic modulus is not infinitely increased but instead has a limit value even on high-stiffness substrates (Mitrossilis et al., 2009; Liu et al., 2013; Trichet et al., 2012).

The strain increment in individual SFs measured from the primary set-point state, *ε*, is specifically described by substituting the force-dependent elastic modulus *E*(*F*) to, for simplicity to capture the main physics, the one-dimensional Hooke’s law

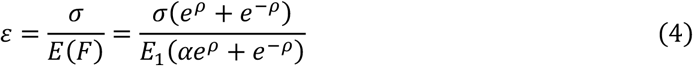

where *σ* is the corresponding stress increment. Differentiating *ε* by *σ* yields

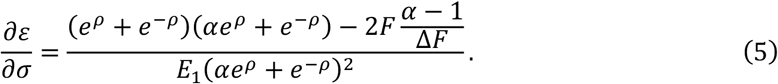

The second derivative of *ε* with respect to *σ* is

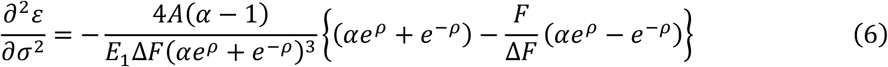

where *A* denotes the cross-sectional area of individual SFs, and hence *F* = *σA. A* may also be modelled using a sigmoid function. However, there is a report that cellular elastic modulus is increased by a factor of several tens (Liu et al., 2018) or at least by 3–4 folds (Liu et al., 2013) upon increased α-SMA expression, while it is unlikely that the cross-sectional area of individual SFs is increased by that amount given that the transition can occur with no substantial change in the lateral width of SF-associated FAs (Goffin et al., 2006). Besides, the critical behavior of this system is determined by a non-dimensional parameter *T*_s_ defined below regardless of the complexity introduced by the second sigmoid function. Thus, we assume *A* is a constant to focus our discussion on the most noticeable observations as well as to make the analysis mathematically more tractable.

At 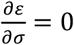 and 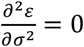 where the slope of the *ε*–*σ* curve is zero at its inflection point, *ρ* = *0*, and hence *F* = *F*_0_; furthermore, in this condition, the following non-dimensional parameter *T*_s_ turns out to be equal to the unity:

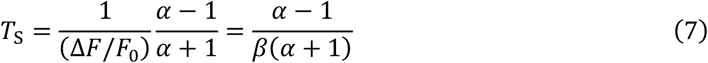

Where

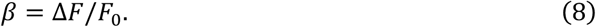

The inflection point of the *ε*–*σ* curve also satisfies

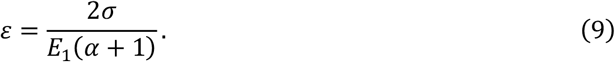

Analysis of the sign of 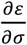 and 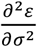 indicates that *ε* always increases with *σ* at *T*_s_ < 1, while *ε* has both a local maximum and a local minimum at *T*_s_ > 1. As described above, this tendency for *T*_s_ to determine the sign of 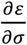 and 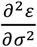 holds also true in a situation where the cross-sectional area *A* is represented as a function of *F* using a sigmoid function similar to that of *E*(*F*). It turns out that strain energy

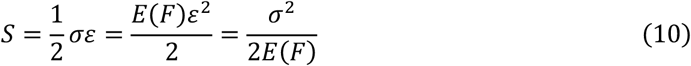

has, as is the case with conventional materials, a single local minimum for stress at *T*_s_ ≤ 1; meanwhile, *S* has two local minima for stress at *T*_s_ > 1.

### 2.1 Parameters

To visualize the results where *ε* and *S* are evaluated as a function of the variable *σ, E*_1_ is assumed to be 5 MPa (Deguchi et al., 2006). *F*_0_ and *A* are assumed to be 10 nN and 1 μm^2^ in accordance with the magnitude of tension in and the cross-sectional area of single SFs, respectively (Deguchi et al., 2006; Matsui et al., 2009). Regarding the rationale for the value of *α*, which measures the ratio of *E*_2_ to *E*_1_ and is defined to be equal to or more than the unity (Fig. 1b), a 3–4 times increase in cellular elastic modulus was detected upon increased α-SMA expression along the length of SFs in valve interstitial cells (Liu et al., 2013). In a separate study, the elastic modulus was increased ∼52-fold (from 0.85 kPa to 44 kPa) for normal human lamina cribrosa cells and ∼30-fold (from 1.1 kPa to 33 kPa) for glaucoma lamina cribrosa cells upon increased α-SMA expression along SFs (Liu et al., 2018). To cover these ranges, an *α* of 1–50 is analyzed. Regarding the value of Δ*F*, Eq. (7) suggests that the relative value of Δ*F*/*F*_0_ (= *β*) essentially matters for the critical parameter *T*_s_ rather than their respective absolute values; thus, *β* is varied under the fixed *F*_0_ value of 10 nN, which then uniquely determines the value of Δ*F*.

## 3. Results

The effect of changing *α* on the relationship between the strain *ε* and stress *σ*, both measured from the primary set-point state, was investigated at a fixed *β* value of 0.2 (Fig. 2a). As mathematically shown above, specific examples illustrate that *ε* increases monotonically with *σ* at *T*_s_ < 1 (specifically, *T*_s_ = 0.0249 with *α* = 1.01). In contrast, *ε* has both a local maximum and a local minimum at *T*_s_ > 1 (with *α* = 5) with an inflection point at *ρ* = 0, i.e., *σ* = *F*_0_/*A* = 10 nN/μm^2^. At the threshold value of *T*_s_ = 1 (with *α* = 1.5), *ε* increases with *σ* except for *σ* = 10 nN/μm^2^ (i.e, *ρ* = 0) where its slope is zero. Accordingly, the strain energy *S* has a single local minimum at *T*_s_ ≤ 1, whereas two local minima appear at *T*_s_ > 1 (Fig. 2b).

**Fig. 2.**
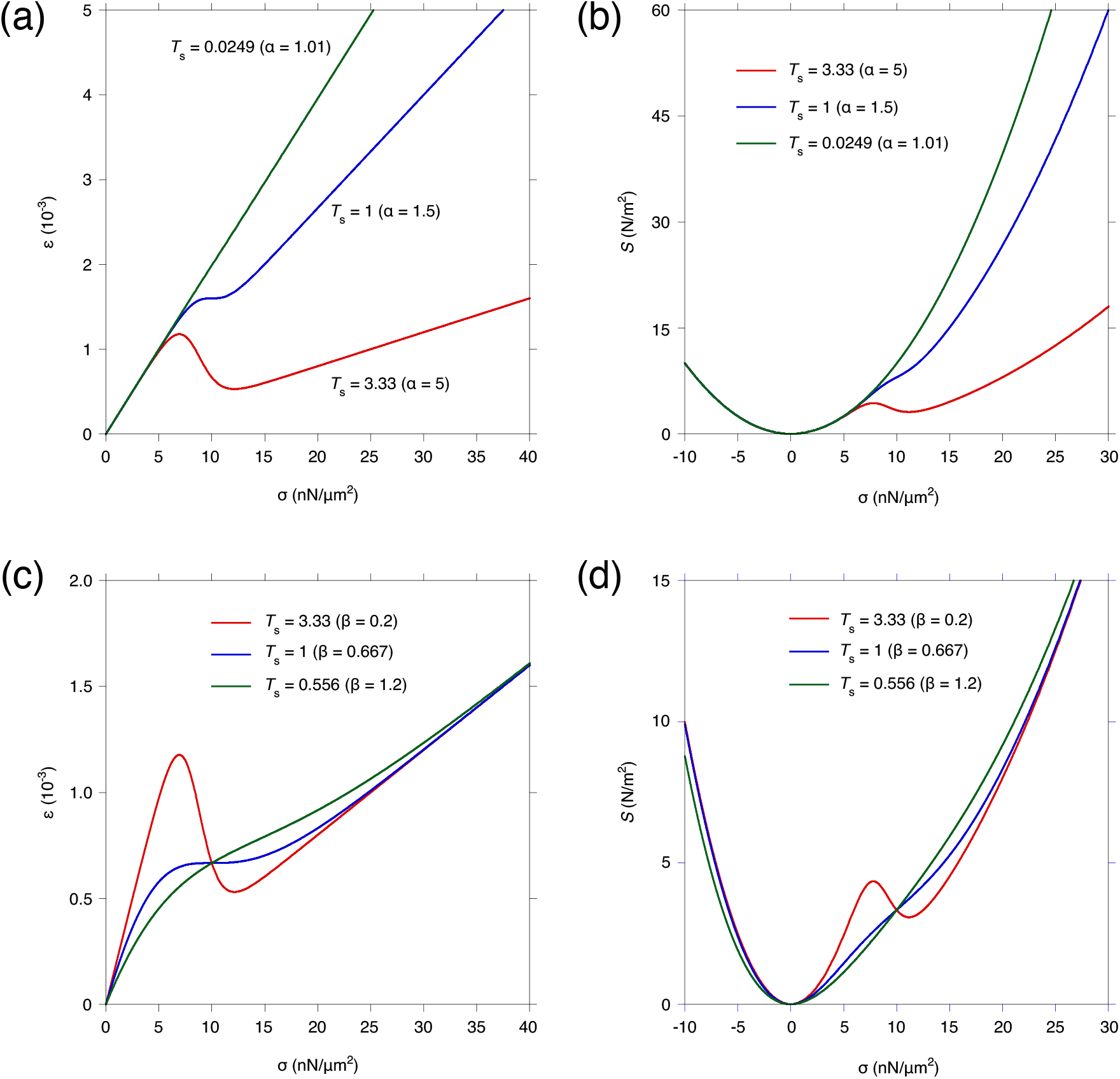
The effect of changing the non-dimensional parameters *T*_s_, *α*, and *β* on the strain *ε* and strain energy *S* as a function of stress *σ*, the variables of which are all measured from the primary set-point state. (a, b) *ε* (a) and *S* (b) with different values of *T*_s_ and *α* at *β* = 0.2. (c, d) *ε* (c) and *S* (d) with different values of *T*_s_ and *β* at *α* = 5.

The effect of changing *β* on the relationship between *ε* and *σ* was likewise investigated at a fixed *α* value of 5. Specific examples demonstrate that *ε* increases monotonically with *σ* at *T*_s_ < 1 (specifically, *T*_s_ = 0.556 with *β* = 1.2). At the threshold value of *T*_s_ = 1 (with *β* = 0.667), *ε* increases with *σ* while having a zero-slope at *σ* = 10 nN/μm^2^ (i.e., *ρ* = 0) (Fig. 2c). As a consequence, *S* has a single local minimum at *T*_s_ ≤ 1, whereas two local minima appear at *T*_s_ > 1 (Fig. 2d). Thus, the emergence of the two stable stress values, corresponding to the two local minima, is observed in a range of *T*_s_ > 1. The inflection point of the *ε*–*σ* curve is independent of *β* as described by Eq. (9), and thus all the curves pass through it (Fig. 2c). Meanwhile, *ε* value at the inflection point is decreased with increasing *α* as is clear from Eq. (9) (Fig. 2a).

Visualization of the comprehensive parametric change in *α* and *β* shows that *T*_s_ > 1, in which two stable stresses are emerged, is achieved at a high and a low value of *α* and *β*, respectively (Fig. 3). In other words, SFs tend to acquire secondary stable stresses if they are subject to as large a change in elastic modulus as possible or to as immediate a transition as possible upon increasing the tension according to Eq. (1).

**Fig. 3.**
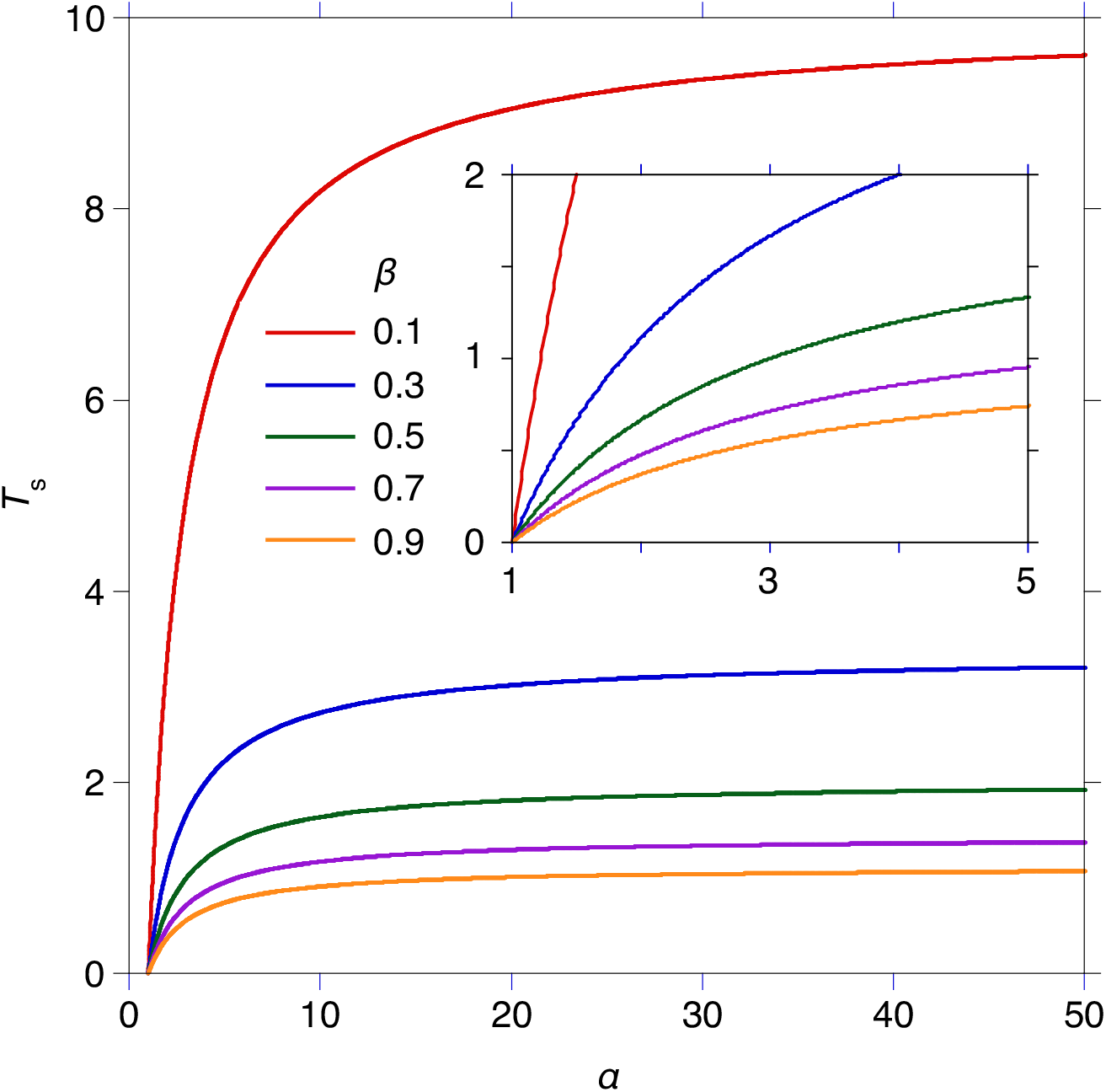
The relationship among the non-dimensional parameters *T*_s_, *α*, and *β*. The inset shows an enlarged view of the same relationship. *T*_s_ = 1 represents the threshold for displaying a secondary stable stress.

## 3. Discussion

The size of individual FAs reversibly increases or decreases according to the tension borne in the connected SFs, eventually allowing the intracellular stress to remain constant over time (Geiger and Bershadsky, 2001; Kaunas and Deguchi, 2011). Meanwhile, if the substrate is stiff enough to bear a large tension in the FA–SF complex, a phase transition occurs to upgrade the homeostatic stress by modulating the molecular composition of the SFs (Goffin et al., 2006). Upon this transition, α-SMA is recruited to such SFs bearing a large tension, leading to an increase in the elastic modulus (Liu et al., 2013; Sakamoto et al., 2016; Liu et al., 2018; Stylianou et al., 2018; Wang et al., 2019). The possession of these homeostatic tensional stresses, known as “tensional homeostasis,” is widely observed across different mesenchymal cell types such as fibroblasts and endothelial cells (Na et al., 2007; Humphrey, 2008; Lu et al., 2008; Deguchi and Sato, 2009; Stamenović and Smith, 2020). Owing to this homeostatic control, cellular tension is relaxed to the basal level even under mechanical stresses such as cell stretch and fluid shear stress (Galbraith et al., 1998; Chien, 2007; Kaunas et al., 2006; Huang et al., 2021). These stresses are known to trigger signaling pathways associated with cell growth and pro-inflammation, while importantly such signals are suppressed over time concomitantly with the recovery of cellular tension. Tensional homeostasis thus allows cells to circumvent chronic activation of pro-inflammatory signals, and its malfunction has been implicated in various diseases that include atherosclerosis and cancer (Hahn and Schwartz, 2009; Kaunas and Deguchi, 2011).

Despite the tremendous significance in biology and medicine, mechanisms of tensional homeostasis remain largely unknown probably because of the necessity of requiring interdisciplinary knowledge related to physics as well as molecular biology. At least, no clear explanation has been put forward regarding the physical origin of the environmental cue-dependent induction of a secondary homeostatic tensional stress. In this regard, our model described here is simple and minimal but capable for the first time of explaining all the key observations in a physically plausible manner. Specifically, it is known that the actual SFs undergo an increased tension upon increasing cellular adhesive area and/or stiffness of the substrate as these two mechanical cues enable efficient tension transmission between the substrate and SFs (Hinz, 2010). Such an increased tension renders the existing SFs less stable, resulting in a phase transition from a β-cytoplasmic actin-dominant phase to an α-SMA-dominant one followed by the stiffening of the updated SFs (Liu et al., 2013; Mitrossilis et al., 2009). The constitutive relation given by Eq. (1) captures this feature. We thereby show that it gives rise to the transition to a state where another homeostatic stress is stably sustained, revealing the physical essence of why multiple set-points exist in cellular tensional homeostasis. Rigorous experimental validation of our theoretical prediction will be the subject of future investigation, but at least it is consistent with existing observations.

Context-dependent protein recruitment to existing SFs is observed not only for α-SMA but also for the septin cytoskeleton (Kang et al., 2020, 2021) and eEF2 elongation factor (Liu et al., 2022) elicited during the epithelial–mesenchymal transition and the cellular senescence, respectively. Enhancement of cellular contractility is followed in these transitions, implying concomitant increase in their elastic properties as well. Given the similarity to the behavior analyzed here, our explanation of why cells end up experiencing different stable stresses may govern such transitions of SFs in general. The present study may also provide insights into why different higher-order forms are present in force-bearing actin-based structures such as podosomes and invadosomes, which are known to take different architectures in a context-dependent manner (Linder et al., 2011; van den Dries et al., 2019). Our theoretical work may thus offer a promising route to addressing such unsolved challenges of highly complicated cellular mechanobiological processes.

## Declaration of competing interest

There are no conflicts of interest to disclose.

## Acknowledgments

This study was supported in part by KAKENHI grant (18H03518 and 21H03796).

## Notes

### Competing Interest Statement

The authors have declared no competing interest.

